# Kidney epithelial proliferation impairs cell viability via energy depletion

**DOI:** 10.1101/2020.04.24.054643

**Authors:** Pierre Galichon, Morgane Lannoy, Li Li, Sophie Vandermeersch, David Legouis, Michael T Valerius, Juliette Hadchouel, Joseph V Bonventre

## Abstract

Both proliferative and anti-proliferative pathways are induced after acute kidney injury. The consequences of proliferation on energy homeostasis and cell viability are unknown. We hypothesized that proliferation regulation is an important determinant of epithelial fate after acute kidney injury. We studied the relationship between proliferation and cell viability in kidney tubular cells. We then analyzed the effect of proliferation on the intracellular ATP/ADP ratio. Finally, we used transcriptomic data from transplanted kidneys to study the relationship between cell proliferation and energy production with different clinical evolutions. We found that proliferation is associated with decreased survival after toxic or energetic stresses in kidney proximal tubular cells. *In vitro*, we found that the ATP/ADP ratio oscillates reproducibly throughout the cell cycle, and that proliferation is instrumental to an overall decrease in intracellular ATP/ADP ratio. *In vivo*, in injured kidneys, we found that proliferation was strongly associated with a specific decrease in the expression of the mitochondria-encoded genes of the oxidative phosphorylation pathway as opposed to the nucleus-encoded ones. These observations suggest that mitochondrial function is the limiting factor for energy production in kidney cells proliferating after injury. The association of increased proliferation and decreased mitochondrial function was associated with poor renal outcomes. In summary, we show that proliferation is an energy demanding process impairing the cellular ability to cope with a toxic or ischemic injury, identifying the association of proliferative repair and metabolic recovery as indispensable and interdependent features for successful kidney repair.

**One Sentence Summary:** Proliferation decreases energy availability in kidney epithelial cells and is associated with enhanced cell death and chronic kidney disease in case of a superimposed metabolic stress.

## INTRODUCTION

Life can be defined as a self-sustained limitation of entropy (negentropy) (*1*). Proliferation is often included in the definitions of life as it pertains both to sustaining a life form and to limiting entropy by increasing the number of energy-containing entities. As such, proliferation - a process indispensable for kidney repair after an acute injury (*2*) - requires energy. Proliferation is tightly controlled under normal conditions and is the cornerstone of development in multicellular organisms. In the presence of various stresses, however, in both prokaryotes or multicellular organisms, pathways are triggered to inhibit proliferation, suggesting that uncontrolled proliferation can be deleterious, even in non-cancerous diseases (*3, 4*). Indeed, the uncontrolled proliferation of repairing kidney tubular cells can lead to chronic kidney disease in case of unrepairable DNA damage and cell cycle arrest (*5, 6*).

Adenosine triphosphate (ATP) is the main energy source for intracellular processes, through its cleavage into adenosine diphosphate (ADP) and hydrogen phosphate (*7, 8*). The intracellular ATP/ADP or ATP/AMP ratio is a more reliable marker of the cell’s energy status than the absolute ATP concentration (*9, 10*). Genetically encoded markers of the ATP/ADP ratio have been developed, allowing its intracellular monitoring in live cells, thus providing a reliable marker of the cell’s energy status (*11, 12*).

A high cellular energy level is generally accepted as a marker of viability (*13*) and is considered a prerequisite for cell proliferation (*14, 15*). Conversely, a persistent decrease in the level of intracellular ATP is associated with cell injury and ultimately death, both in tumor and non-tumor cells (*16, 17*). It is generally assumed that the ATP/ADP ratio is regulated prior to changes in proliferation (*18–21*). Indeed, a low ATP/ADP ratio (or its close correlate, ATP/AMP) activates AMPK, which induces cell cycle arrest through P27, P53 and P21 activation in various organs (*22*). By contrast, in cancer, a low ATP/ADP ratio is considered to allow cell proliferation by stimulating aerobic glycolysis and the Warburg effect (*10*). Although the effect of ATP/ADP variations on proliferation has been studied, the reverse mechanism, i.e. the effect of cell cycle progression on the cellular energy status, is yet unexplored. Because the ATP/ADP ratio reflects the energy in a cell (*23–25*), and energy depletion leads to cell death (*17, 26*), we hypothesized that the effect of proliferation on the ATP/ADP ratio might impact the cellular fate more strongly than the regulatory feedback on proliferation triggered by changes in the ATP/ADP ratio. Here, using viability assays and transcriptomic analyses, we investigated the relationship between proliferation and viability in human cancer and renal epithelial cells. By quantifying the ATP/ADP ratio in live cells using a single cell approach, we studied the effect of proliferation on the intracellular ATP/ADP ratio, and characterize the pattern of variation of the intracellular energy level throughout the cell cycle. Lastly, we investigate the differential transcriptomic signature of energy production associated with proliferation in kidney allografts after an ischemic injury, identifying mitochondrial-encoded genes of the oxidative phosphorylation pathway as a limiting factor associated with proliferation in kidneys transitioning to chronic kidney disease after an episode of acute kidney injury.

## RESULTS

### Proliferating cells are more sensitive to an injury than non-proliferating cells

We subjected human renal epithelial cells (HK2 cell line) to a toxic or metabolic stress when their proliferation was either stimulated by pifithrin-alpha (a P53 antagonist promoting G1/S and G2/M transition) or inhibited by tenovin-1 (a P53 agonist inhibiting the G1/S transition) (Fig. 1A-B). We observed that the stimulation of cell proliferation by pifithrin-alpha promoted the death of the cells exposed to puromycin (a well characterized toxicant acting through inhibition of protein synthesis), whereas inhibition of cell proliferation by tenovin-1 was protective. Pifithrin-alpha alone or tenovin-1 alone was not toxic, as they did not cause cell death in the absence of puromycin (Fig. 1A). Similar results were obtained when cells were cultured under conditions of energy depletion (no oxygen, no glucose) versus control conditions (normoxia, glucose) (Fig. 1B). Taken together, these data show that proliferation enhances susceptibility to noxious stimuli or impairs the ability of cells to survive an injury.

**Fig. 1:**
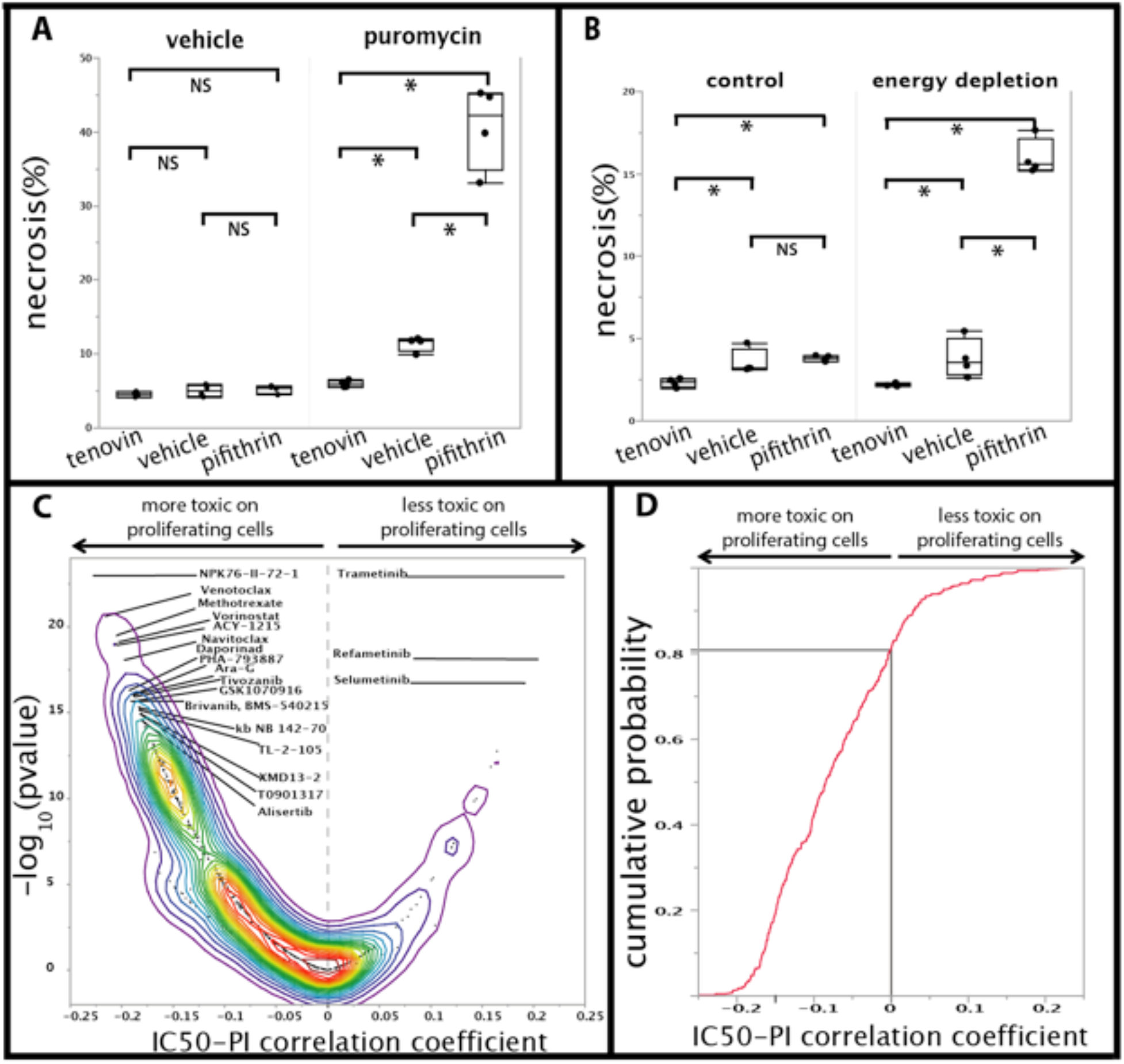
Consequences of cell proliferation on susceptibility to cell death. (**A**). Effect of cell cycle inhibition by tenovin-1 or cell cycle facilitation by pifithrin-α on cellular death caused by puromycin. (**B**). Effect of cell cycle inhibition by tenovin-1 or cell cycle facilitation by pifithrin-α on cellular necrosis caused by energy depletion (oxygen and glucose deprivation). (**C**). Vulcano plot of the correlation of the cytotoxicity index with the proliferation index tested for 367 compounds. The top 20 significant correlations are labeled. (**D**). Cumulative probability for all 367 compounds is plotted against the coefficient of the correlation between IC50 and proliferation index, showing a skewed distribution towards correlation between proliferation and chemosensitivity. *: p-value <0.05; NS: statistically nonsignificant.

In order to test this hypothesis in a large number of different cell types, we used the Genomic of Drug Sensitivity in Cancer (GDSC) database, where the chemosensitivity of 1021 different cancer-derived cell lines is measured for 367 different molecules (*27*). We found a significant correlation (*P*<0.05) of chemosensitivity with the proliferation index for 257 molecules (70% [65-75], *P*=1.1e-14). The correlation was positive for 90% [86-94] of these molecules (*P*=2.2e-16) (Fig. 1C-D). Interestingly, chemosensitivity was most associated to proliferation for drugs targeting cell cycle, metabolism and apoptosis, whereas chemoresistance was most associated with proliferation for MEK1/2 inhibitors (Table S1).

Given that proliferation requires energy and that a decreased cellular energy level results in cell death, we hypothesized that the depletion of cellular energy in proliferating cells could be responsible for this enhanced susceptibility to toxic influences.

### The ATP/ADP ratio decreases when cells proliferate

In order to monitor the cellular energy level during the course of a cell cycle, we measured the ATP/ADP ratio in proliferating human renal epithelial cells (HK2 cell line). PercevalHR is a genetically encoded fluorescent marker that enables the live monitoring of the ATP/ADP ratio(*11, 12*). To monitor the single cell-ATP/ADP ratio over time in proliferating epithelial cells, we generated human renal epithelial HK2 cells stably expressing PercevalHR (Fig. S1A). We first validated this quantification of the ATP/ADP ratio (ATP/ADP[Perceval]) by an independent technique based on the measurement of luciferin/luciferase bioluminescence (ATP/ADP[Luciferin]). The total ATP/ADP[Perceval] signal correlated well with the ATP/ADP[Luciferin] (r=0.96, p=0.0001). We also observed the expected decrease of ATP/ADP[Perceval] after chemical energy depletion with blockage of glycolysis and oxidative metabolism using 2-deoxyglucose (2 DG) and sodium azide (NaN_3_) (Fig. S1B-C).

Single cell ATP/ADP[Perceval] was quantitated in real-time *in vitro*. Although changes in the mean ATP/ADP values over time were reproducible under basal conditions or during energy depletion, this ratio was highly variable among proliferating cells in the same well and at the same time (Fig. S1C). This observation suggests that the ATP/ADP ratio is a dynamic parameter, undergoing variations that are at least partly independent of the extracellular environment. We hypothesized that the ATP/ADP ratio could be correlated with the cell cycle stage. To test this hypothesis, we plated PercevalHR-expressing epithelial cells at various densities to achieve different levels of contact inhibition of proliferation (Fig. 2A, left panel). We found that the average intracellular ATP/ADP ratio increased when the cells were more confluent. To rule out a direct effect of the confluency, we cultured the cells at the same densities in the presence of rigosertib, a potent polo-like-kinase 1 (PLK1) inhibitor inducing G2/M arrest. Non-proliferating cells showed a significant increase in the ATP/ADP ratio, regardless of their degree of confluence. Conversely, performing a scratch assay on confluent cells induced the proliferation and migration of the cells, which then displayed a low ATP/ADP ratio (Fig. 2A, right panel). Again, rigosertib increased the cellular ATP/ADP ratio while inhibiting wound closure.

**Fig. 2:**
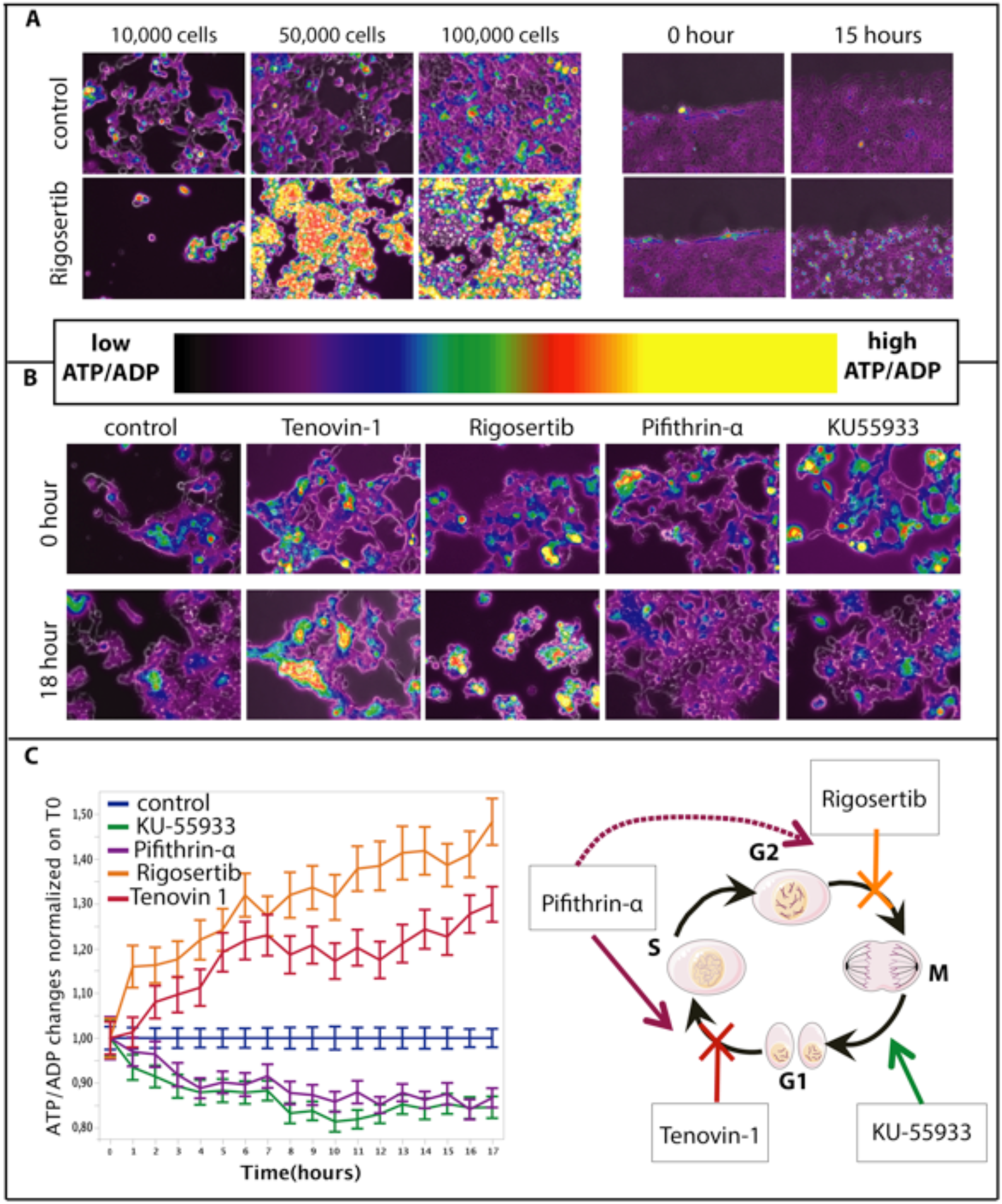
Cell cycle interventions and their effects on ATP/ADP ratio. **(A). Left panel:** Wells were plated with increasing numbers of PercevalHR expressing cells to obtain various levels of confluence in control condition or with 1μM rigosertib to inhibit the cell cycle. **Right panel:** Representative pictures obtained 0 and 15 hours after a scratch performed under control condition or with exposure to 1μM rigosertib to inhibit wound closure. (**B**). Pharmaceutical interventions on the cell cycle using various compounds on PercevalHR expressing cells over an 18-hr time course: KU55933 (10 μM), pifithrin-α (50 μM), rigosertib (10 μM), tenovin-1 (10 μM). **c**. Quantification of changes in single cell intracellular ATP/ADP ratio over an 18-hr exposure to various cell cycle modifiers.

To exclude a possible cell cycle-independent effect of rigosertib, we then used a panel of pharmaceutical compounds to study the changes in the ATP/ADP ratio over time when the cell cycle is perturbed (Fig. 2B-C). Tenovin-1 and Rigosertib were used to inhibit the cell cycle during the G1/S or G2/M transition, respectively. Conversely, pifithrin-alpha and KU-55933 (an ATM inhibitor promoting G2/M transition) were used to stimulate cell proliferation. Single-cell monitoring demonstrated that cell cycle inhibition increased ATP/ADP progressively over time whereas cell proliferation decreased it (Fig. 2C).

### Cell cycle and proliferation-dependent variations in the intracellular ATP/ADP ratio

The single-cell ATP/ADP ratio was maximal around cytokinesis in non-synchronized cells (Fig. 3A, left panel). Variations of the ATP/ADP ratio before and after cytokinesis follow a systematic pattern (Fig. 3A, right panel): after a gradual increase, the ratio peaks at mitosis before dropping dramatically. This pattern was highly reproducible, including in cells exposed to pharmaceutical inhibition or stimulation of the cell cycle (Fig. 3b). However, the cell cycle inhibitor tenovin-1 increased the mean ATP/ADP ratio over time compared to the cell cycle facilitator pifithrin-alpha(Fig. 3B). Thus, the intracellular energy level, evaluated by the ATP/ADP ratio, is directly influenced by the cell cycle.

**Fig. 3:**
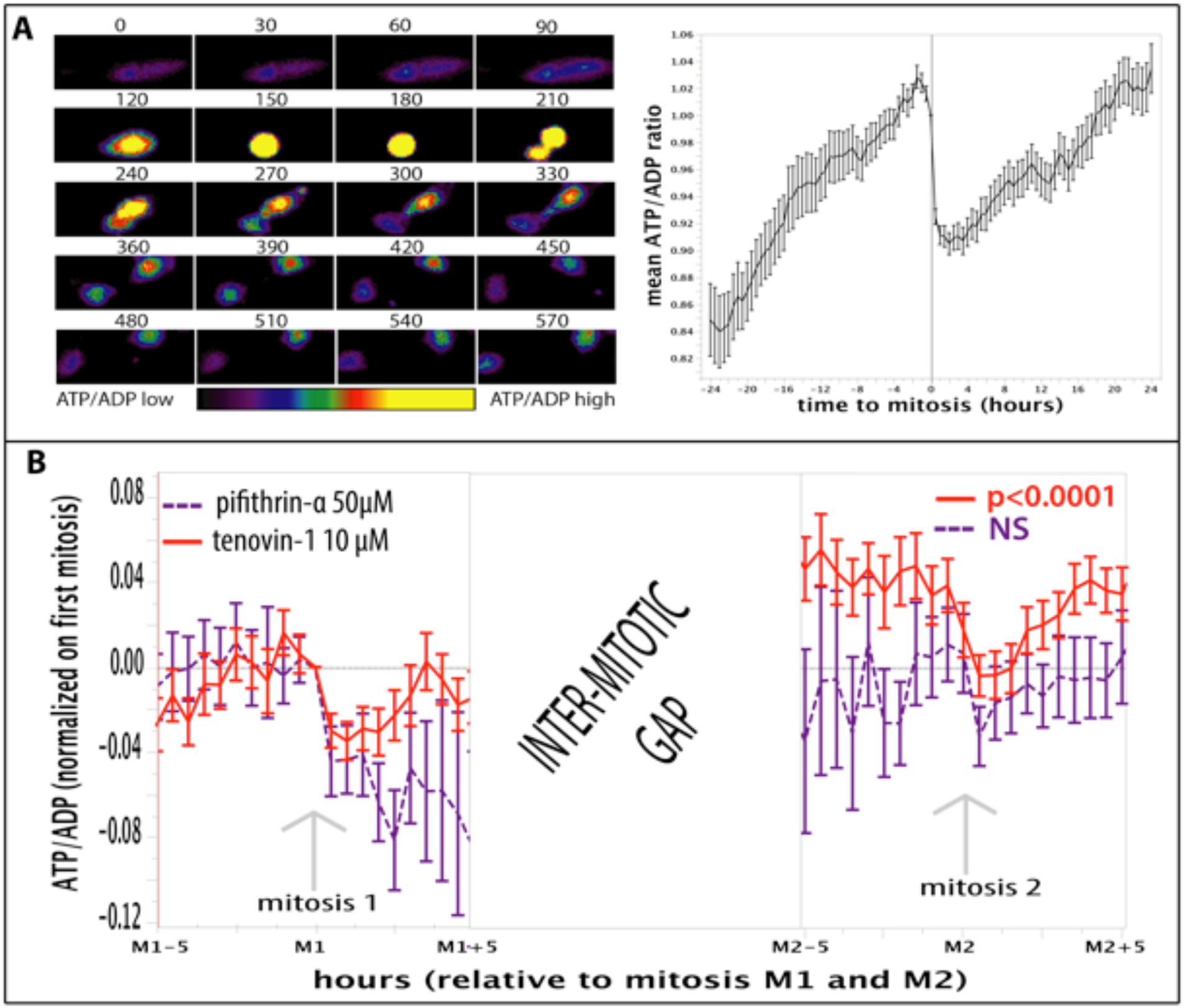
ATP/ADP variations throughout the cell cycle. (**A**, left panel): Live imaging on a single PercevalHR expressing cell undergoing mitosis. (**A**, right panel): Quantification of the intracellular ATP/ADP ratio in single cells during the perimitotic period with *in silico* synchronization on time of mitosis. **(B)**. ATP/ADP changes in PercevalHR expressing cells incubated with the cell cycle facilitator pifithrin-α or with the cell cycle inhibitor tenovin-1 between 2 successive mitoses. ATP/ADP ratio values are represented as means and whiskers indicate the standard error. P-values represent the significance of the comparison between the first and second mitosis within each treatment group, using a Wilcoxon rank test.

In order to study potential variations in the single cell ATP/ADP ratio in successive cell cycles within the same cell, ATP/ADP ratio trajectories were evaluated in cells undergoing division twice in the same live-imaging experiment. The cells had the same pattern of ATP/ADP variations for each of the two successive mitoses. After a first mitosis, cells treated with tenovin-1 reached the second mitosis with a significantly higher ATP/ADP ratio than at the first mitosis, compared to cells treated with pifithrin-alpha in which no such an increment occured (Fig. 3B). We conclude that cell cycle inhibition increases ATP/ADP ratio in cells independently of the cell cycle stage, in a timedependent manner: the slower the cell cycles, the more positive the cellular energy balance.

### Association of a mitochondrial defect with proliferation in injured kidney allografts

To test the relevance of our findings in proliferating cells *in vivo*, we studied the transcriptomic profiles of 8375 tumors from The Cancer Genome Atlas (TCGA) database. We computed an estimate of the proliferation index, using the median value of a set of proliferation-associated genes, as previously described (*28*). Pathway analysis was based on the correlation of each gene expression profile with the computed proliferation index, with focus on the three energy-producing pathways from the Kyoto Encyclopedia of Genes and Genomes (KEGG): oxidative phosphorylation, citrate cycle, and glycolysis/neoglucogenesis (*29*). We found that the level of expression of the genes involved in ATP generation (oxidative phosphorylation or OXPHOS) was positively correlated with proliferation in cancers.

We then investigated the relationship between proliferation and ATP generation in injured kidneys by analyzing the data from kidney allografts before and after implantation (*30*). We found proliferation and OXPHOS to be negatively correlated in kidney allografts after ischemia reperfusion (Fig. 4A and Tables S2 and S3).

**Fig. 4.**
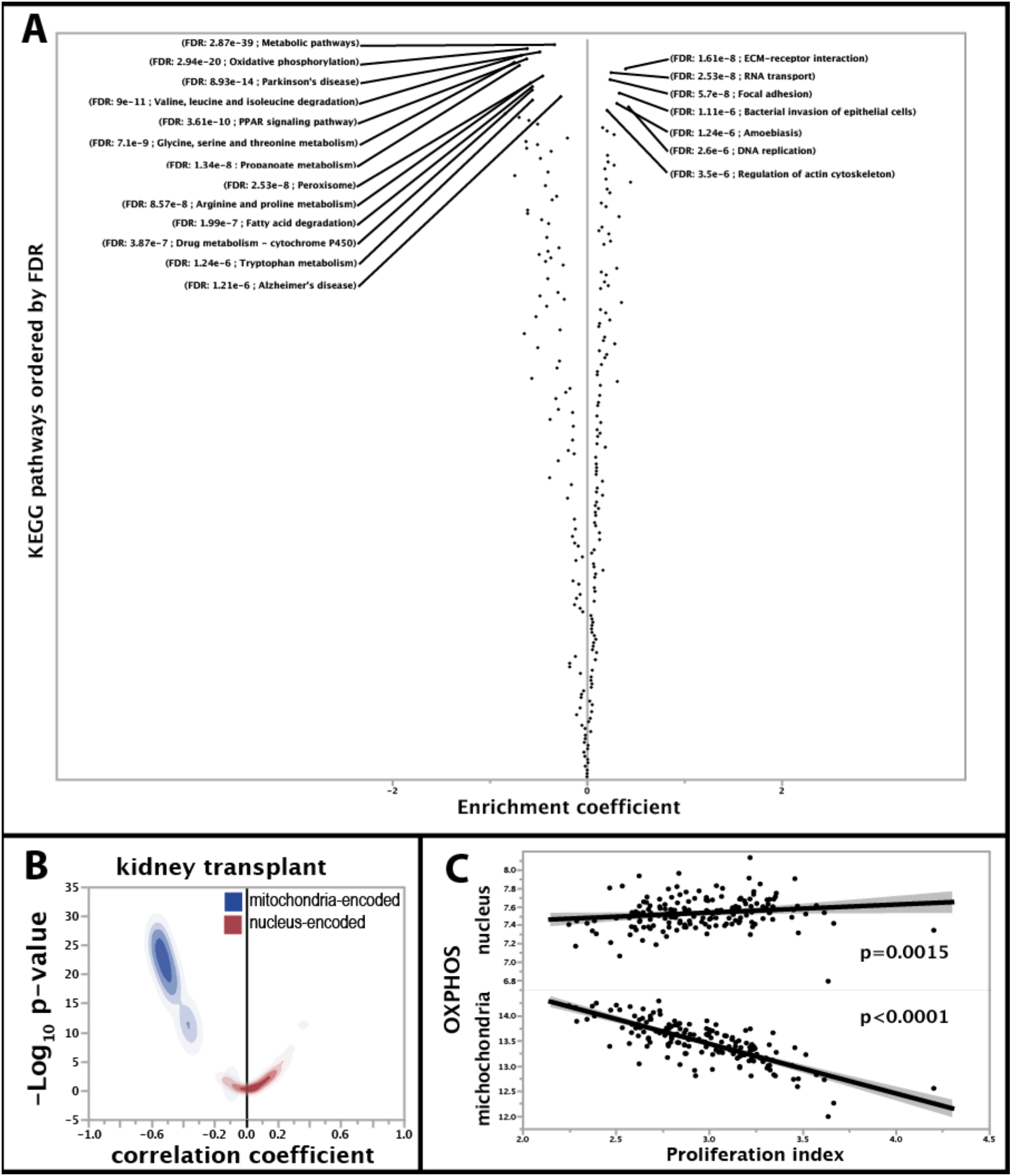
Transcriptomic signature associated with proliferation. **(A)**. KEGG pathways enrichment associated with proliferation in kidney allografts immediately after reperfusion. The 20 most significant pathways are labeled. **(B)**. Vulcano plot showing the correlation between OXPHOS gene expression and the computed proliferation index in kidney allografts (right panel). x-axis: Kendall’s k correlation coefficient; y-axis: significance represented by the −log(p-value) of the correlation (Kendall); blue: mitochondria-encoded genes; red: nucleus-encoded genes. **(C)**. Differential correlations of nucleus and mitochondria-encoded OXPHOS with the proliferation index in kidney allografts.

A further analysis showed that the decrease in OXPHOS genes expression with proliferation in kidney allografts was the result of a profound downregulation of the expression of mitochondria-encoded genes. In contrast, the transcription of the nucleus-encoded OXPHOS genes was moderately upregulated with proliferation (Fig. 4B-C and Fig. S2). Using single-cell RNA-seq data from rejecting human kidneys (*31*), we analyzed the relationship between proliferation and OXPHOS related genes specifically in renal epithelial cells (Fig. 5A and S3). This new analysis confirmed that proliferation was negatively correlated with mitochondria-encoded OXPHOS genes, but not with nucleus-encoded OXPHOS genes (Fig. 5B).

**Fig. 5.**
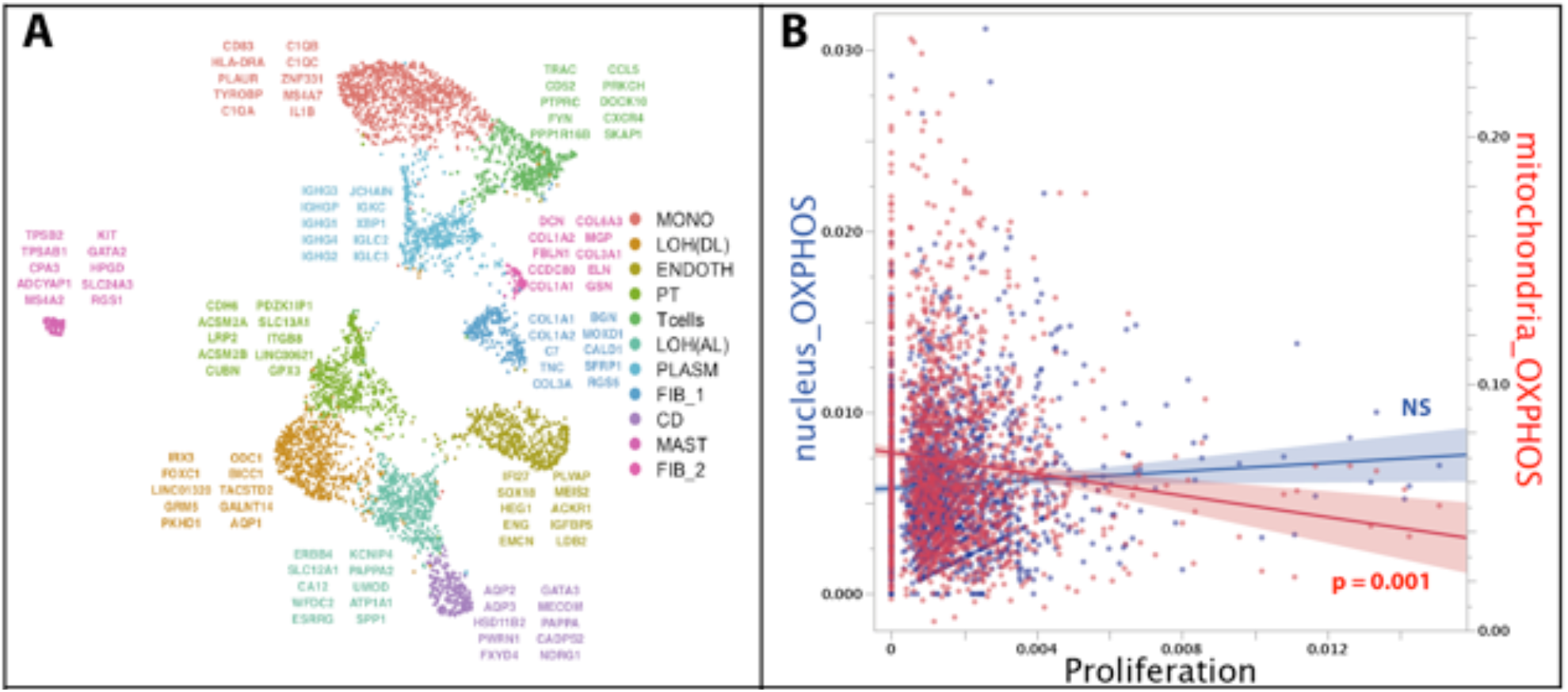
Oxidative phosphorylation in proliferating epithelial cells. **(A).** Single cell analysis of human rejecting kidney showing clusters corresponding to different cell types within the kidney. **(B).** In the subset of renal epithelial cells (PT, LOH and CD), correlation of proliferation with nucleus or mitochondria-encoded OXPHOS genes.

### Association of proliferation, mitochondrial status and outcomes

The study of Cippà and colleagues was performed on biopsies performed on the same kidney allografts before, just after, 3 and 12 months after transplantation (*30*). The first two series of samples constituted the ‘early’ group, while the last two constituted the ‘late’ group. Furthermore, three types of evolution were distinguished within the late biopsies: recovery, transition towards chronic kidney disease and established chronic kidney disease (CKD) (*30*).

As shown in Fig. 6A, the computed proliferation index was higher in the kidney biopsies performed 3 or 12 months after transplantation than in biopsies performed immediately before or after transplantation. Moreover, a higher computed proliferation index was associated with the progression towards CKD (Fig. 6B). OXPHOS transcripts were found to be downregulated in the nucleus only during the acute phase of injury (reperfusion), whereas they were downregulated in the mitochondria during the transition towards chronic kidney disease (Fig. 6D,E,G,H).

**Fig. 6.**
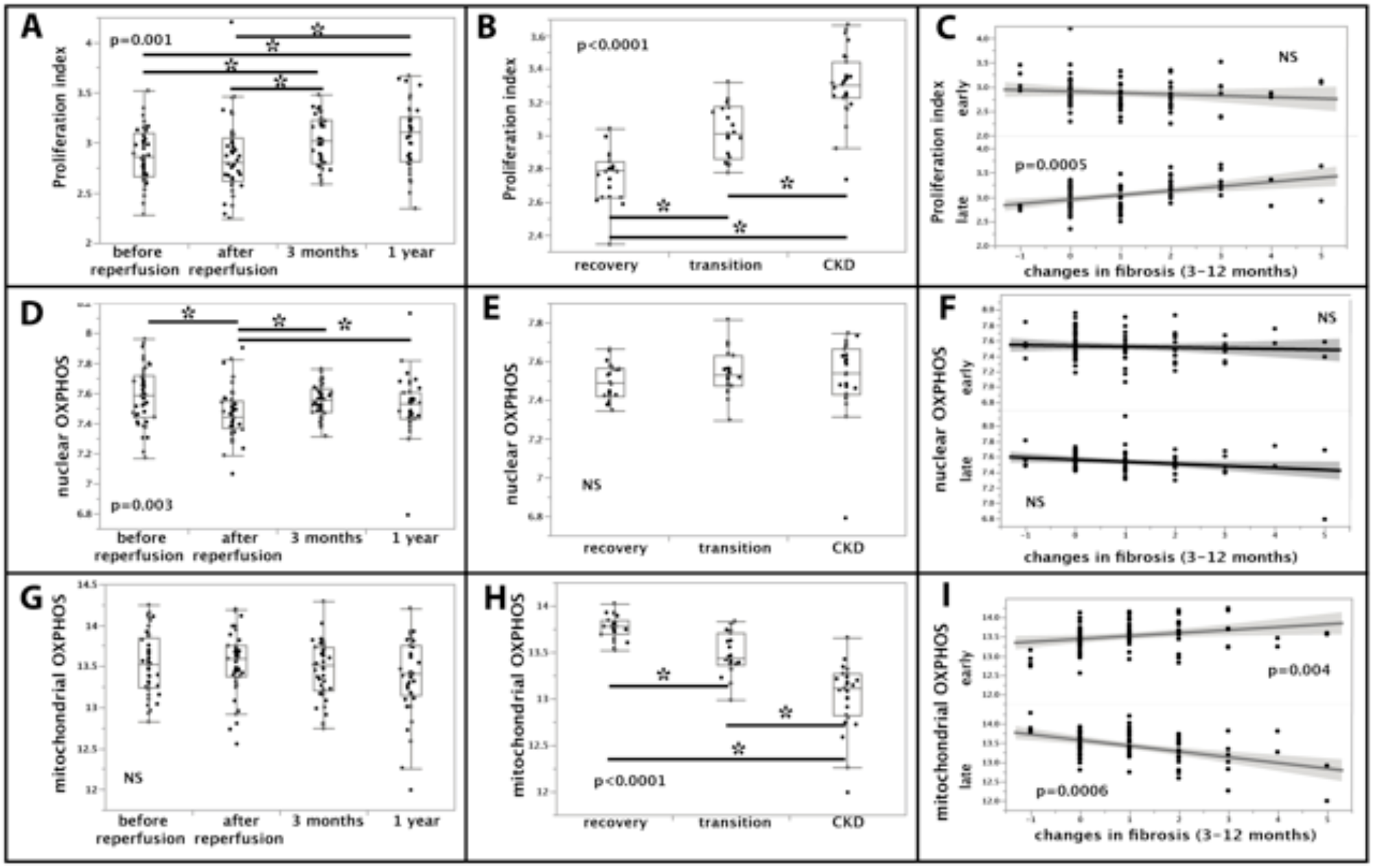
Association of proliferation index and OXPHOS with chronic kidney disease progression. **(A-C).** Proliferation index in respect to different timings, evolution profiles and fibrosis progression. **(D-F).** Nuclear OXPHOS index in respect to different timings, evolution profiles and fibrosis progression. **(G-I).** Mitochondrial OXPHOS index in respect to different timings, evolution profiles and fibrosis progression. *: p-value<0.05 by Wilcoxon test (by pairs), when p-value<0.05 (as indicated on graphs) by Kruskall-Wallis test.

We then studied the relationship between proliferation, the regulation of OXPHOS genes and the long-term clinical outcomes, i.e. the glomerular filtration rate (GFR) and incidence of fibrosis (Fig. 6C,F,I and Supplemental Fig. 4. We found that the timing dramatically modified the relationship between proliferation, the OXPHOS level and the incidence of fibrosis in the kidney. If there was no correlation between the proliferation index and fibrosis in the early biopsies, a low mitochondrial OXPHOS index in the early biopsies (immediately before and after transplantation) was associated with a lesser incidence of fibrosis in the kidney between 3 and 12 months after transplantation (Fig. 6I). In contrast, a decrease in the mitochondrial OXPHOS index and an increase in the proliferation index in the late biopsies (3 and 12 months after transplantation) were associated with an increased fibrosis and a decreased GFR at 12 months (Fig. 6I and Fig. S4).

Taken together, these transcriptomic analyses suggest that the decrease of mitochondria-encoded genes represents a limiting factor for the energy production by OXPHOS required for the proliferative repair of the kidney allografts after injury.

## DISCUSSION

Here we show that proliferating cells are more sensitive to injury than non-proliferating ones, and that the inhibition of proliferation is protective. We show that proliferation causes a decrease in intracellular energy with superimposed energy oscillations depending on the cell-cycle stage. *In vivo*, we found a correlation between proliferation and the expression of nucleus-encoded genes of the OXPHOS gene set, indicating a transcriptomic program oriented towards energy production in proliferating cells. However, mitochondria-encoded genes were profoundly downregulated in the proliferating kidney after injury, indicating a mitochondrial failure to back up the nuclear induction of OXPHOS genes. Therefore, proliferative repair is a state of particular vulnerability for the kidney. Finally, we found that the persistence of proliferation associated with the downregulation of mitochondrial OXPHOS genes is a pattern associated with an unfavorable evolution towards chronic kidney disease.

### Proliferation and energy metabolism are interdependent

The fact that a low intracellular ATP induces molecular pathways downstream of AMPK to control proliferation suggests the presence of a regulatory feedback to mitigate the decrease in intracellular ATP levels caused by increased energy consumption. Such an effect of proliferation on ATP could explain why alleviating cell cycle blockade by P21 or P53 aggravates the course of acute kidney injury caused by an ischemic episode (*2*), and why highly proliferative tumors frequently undergo spontaneous necrosis (*17*). The kidney energy turnover is very high. It is thus very sensitive to injury (and especially ischemia), which causes an immediate decrease in energy production, overall energy depletion and epithelial cell death (*32*). After an injury, cell proliferation is necessary to replace lost cells (*33*). Surprisingly, an increase in cell proliferation caused by the inactivation of the anti-proliferative factors p53 and p21 was shown to worsen the lesions caused by an episode of ischemia-reperfusion in the kidney (*34–36*). The p53 pathway is important to arrest the cell cycle in case of DNA damage, delaying proliferation until DNA is repaired (*6*). DNA repair is a highly energy-demanding process fueled by mitochondrial ATP generation (*37*). Thus, proliferation arrest might be an energy-saving mechanism, especially in an injured state. In agreement with this hypothesis, we found an association between increased proliferation, ATP depletion and increased cell death in renal epithelial cells in which *Nupr1* (Nuclear Protein 1) was inactivated (*38*). Nupr1 is a downstream effector of ATF4, a master regulator of endoplasmic reticulum stress, the activation of which leads to excessive protein synthesis, causing energy depletion and cell death (*39*). Taken together, these studies suggest that an increase in cell proliferation in conditions of stress might cause critical energy depletion due to enhanced energy needs for both cell maintenance and biosynthetic processes, thus resulting in cell death if a critical energy threshold is not maintained (*40*).

### Intercellular and intracellular variations in energy metabolism

Our observations change the understanding of energy variations in proliferating cells (Fig. 7). Previous experiments were limited by the fact that timed bulk cell analysis did not allow the study of the ATP/ADP trajectories of single cells within the same extracellular environment. We show by single-cell analysis that a progressive increase in the ATP/ADP ratio in proliferating cells can be observed if there is increasing confluence and contact inhibition, whereas proliferation itself (at the single-cell level) is associated with decreased ATP/ADP ratio. In addition, we show that within proliferating cells, the ATP/ADP ratio oscillates with the cell cycle. These oscillations in ATP/ADP ratio are reminiscent of cyclic variations of the TCA cycle flux described by Ahn and colleagues (*41*), and this is in keeping with our data *in vivo* showing that OXPHOS transcripts are overexpressed in proliferating cells. The intracellular energy level not only varies during the cell cycle but also depends on the proliferation rate. The live monitoring of the ATP/ADP ratio in single cells demonstrates transient physiological decreases in ATP/ADP in proliferating cells. Although well tolerated under basal conditions, these proliferating cells with lower ATP/ADP ratio were more vulnerable to a superimposed injury than non-proliferating cells.

**Fig. 7.**
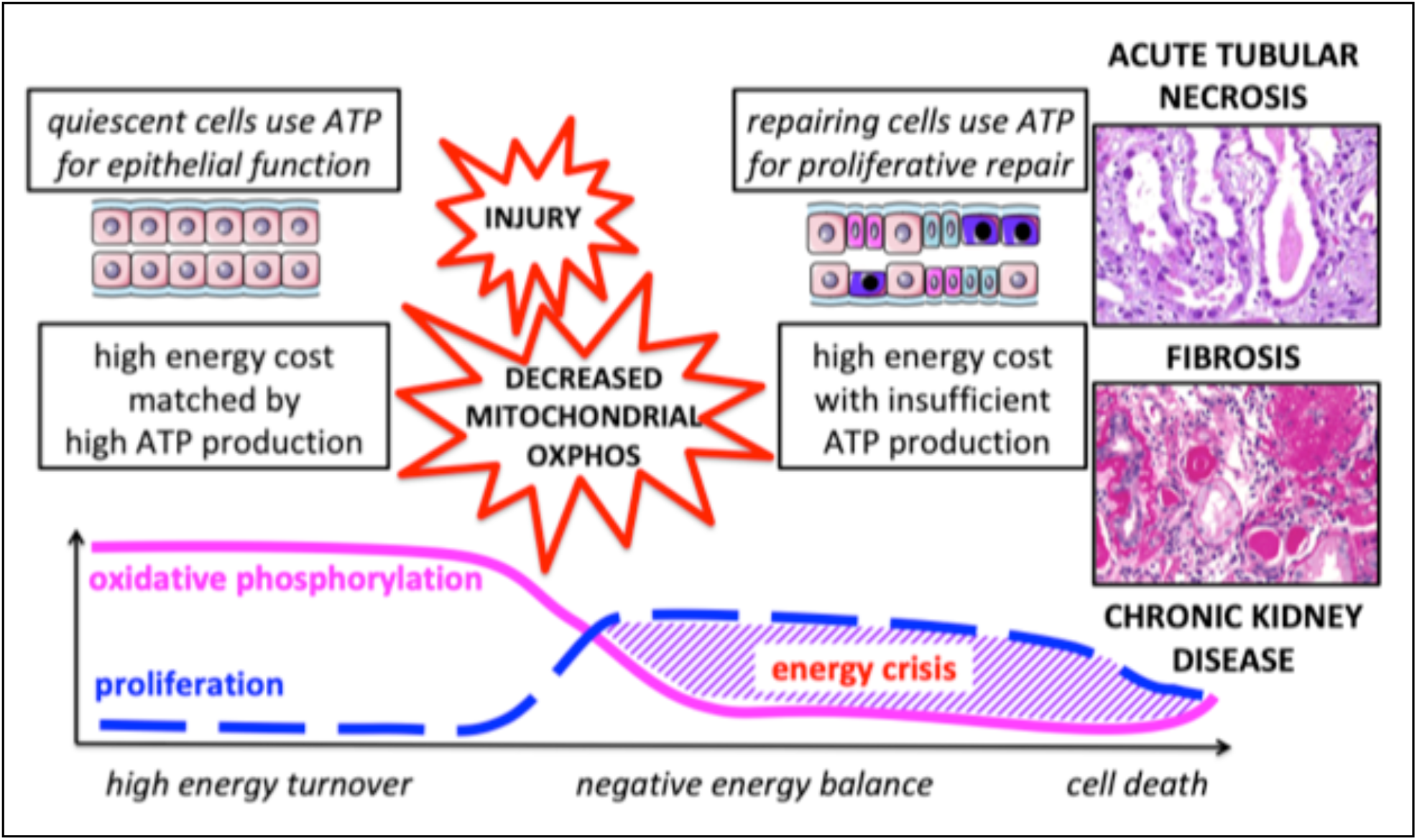
Impact of energy metabolism changes leading to cellular injury in proliferative cancers treated with chemotherapy and in injured kidneys.

Mendelsohn et al. found that cell growth and ATP can follow dissociated patterns in a high throughput CRISPRi screen (*42*). We provide experimental evidence that intracellular energetic levels (ATP/ADP ratio) do not determine cell proliferation. Quite the opposite, we show that proliferation negatively impacts the ATP/ADP ratio. In addition, upon injury, cell survival is improved by the inhibition of proliferation, whereas cells in which proliferation is stimulated are more sensitive to injury. This is likely related to an accelerated energy consumption required for protein synthesis and other metabolic processes necessary for genetic amplification prior to cell division.

### Effect of proliferation on cell viability upon injury

We find proliferation to be associated with chemosensitivity, as previously described (*20, 43–45*) and provide experimental data to demonstrate the causal relationship between proliferation and sensitivity to cell death. Considering that cancer cells with a high intracellular ATP level are more resistant to cytotoxic chemotherapy (*46*), proliferation control might be a mechanism for cancer to resist to chemotherapy or ischemia by maintaining high intracellular ATP/ADP ratio. Similarly, inhibiting proliferation might be a protective mechanism to prevent lethal energy depletion in epithelial cells. This protective mechanism might be responsible for the protective effect of the extensively studied phenomenon of ischemic preconditioning (*47–49*), where a mild and transient ischemia triggers cytoprotective pathways concomitant with transient cell cycle arrest (*50, 51*). Among these pathways, p53, p21 and AMPK were shown to protect the kidneys from ischemiareperfusion injury (*52*) while suppressing cell proliferation (*4, 53*).

However, cell proliferation is absolutely necessary for vital processes. Following a renal epithelial injury, epithelial proliferation is known to be instrumental to organ repair (*33*). Proliferation demands a high energetic input, and oxidative phosphorylation is the most efficient pathway for energy production. In keeping with that, we found that proliferation is associated with increased expression of nucleus-encoded OXPHOS genes. Kidneys with an unfavorable evolution profile had a dissociated transcriptomic pattern (high proliferation index with maintained expression of nucleus-encoded OXPHOS genes but decreased expression of mitochondria-encoded OXPHOS genes). This is in keeping with recent studies showing that maintaining oxidative phosphorylation during and after acute kidney injury (AKI) is protective (*54–56*), and other studies suggesting that energy-parsimonious approaches might mitigate renal damage at the early stage of acute kidney injury. Stabilization of the hypoxia inducible factor (HIF) shifts the energy metabolism to anaerobic glycolysis, inhibits cell proliferation (*3, 57, 58*) and protects against ischemic acute kidney injury (*59–61*).

### Limitations to the study of the effect of proliferation

We found that proliferation is associated with decreased viability of kidney epithelial cells, and that intervention on cell proliferation modifies the ATP/ADP ratio and cell viability. Although this suggests that proliferation causes energy depletion and decreased viability, one cannot rule out the possibility that our interventions on cell proliferation may have independently impacted the energy metabolism and survival pathways. Such off-target effects of cell cycle acting pathways have been described, e.g. for p53. Regulation of the cell cycle is an important mechanism to allow the repair of DNA damage in the injured kidney (*6*). However, our results were consistent both when comparing cells with naturally occurring variations in proliferation rate and with various cell cycle acting agents targeting different checkpoints. In order to further affirm the causal link between proliferation, decrease in ATP/ADP ratio and cell viability, one would need to be able to restore higher levels intracellular ATP/ADP ratio in proliferating cells without causing disturbances in metabolic or survival pathways, which to the best of our knowledge cannot be achieved. Thus, this limitation is rather conceptual and does not affect the conclusions and translational perspectives of our study.

### Proliferation as a therapeutic target in kidney diseases

We identified kidney epithelial proliferation as a state of energy crisis with increased sensitivity to cell death. However, epithelial cell proliferation is necessary for kidney repair. Thus, an interesting strategy to mitigate necrosis would be to delay proliferative repair in kidneys with ongoing injury, and allow proliferation after cessation of the injury.

Identifying the time frame for proliferation inhibition after kidney injury in individual patients will necessitate the implementation of dynamic biomarkers of acute kidney injury. In this perspective, the cell cycle arrest markers IGFB7 and TIMP2 are clinically available urinary markers that might help to refine the different stages of injury and repair after acute kidney injury (*62*). Urinary quinolinate/tryptophan ratio is another emerging non-invasive marker allowing to pinpoint the mitochondrial defect during acute kidney injury (*55*).

Targeting the cell cycle to orchestrate kidney repair is an attractive option, but raises the concern of extrarenal side effects, as proliferation is both necessary for the self-renewal of certain tissues like gut or blood cells, and also instrumental in the development of cancers. Thus, targeting proliferation in a time and tissue specific manner would be an advantageous strategy to improve renal outcomes while minimizing the risk of side effects.

In summary, proliferation is an energy demanding process impairing the cellular ability to cope with a toxic or ischemic injury. We postulate that interventions to mitigate proliferation and restore energy production can enhance cell survival and organ recovery (Fig. 7).

## MATERIALS AND METHODS

### Study design

The starting hypothesis of this study was that proliferation may critically affect the energy homeostasis in human kidney epithelial cells.

We first tested our hypothesis *in vitro*, by increasing or decreasing proliferation rates in kidney epithelial cells with various methods and assessing cell viability and intracellular ATP/ADP ratio at the single cell level. The generalizability of our results was tested on the large cytotoxicity assay of the Genomic of Drug Sensitivity in Cancer (GDSC) database, testing 367 different molecules in 1021 different cancer-derived cell lines(*27*).

We then used RNA-seq data from kidney allograft biopsies in order to test the relationship between proliferation and energy metabolism *in vivo* in 163 kidney allograft biopsies from 42 patients (*30*). Single cell RNA-seq data was used to confirm that the results were specifically applicable to kidney epithelial cells (*31*). A large dataset of 8375 tumors from The Cancer Genome Atlas (TCGA) database (*63*) was used to assess the generalizability of our results, and identify the specificities of injured kidneys compared to various growing tissues.

Lastly, we studied the clinical correspondences of cell proliferation and energy metabolism in injured kidneys in terms of recovery or progression to chronic kidney disease in 42 kidney allograft patients(*30*).

### Cell cultures

HK2 (ATCC^®^ CRL-2190™) cells are immortalized male adult human renal proximal tubular cells cultured at 37°C in DMEM with 10% fetal bovine serum. The live microscopy experiments were conducted in Leibovitz’s L-15 medium with no phenol red (Fisher, #21083027). L-15 was specifically designed for renal primary cell cultures. It provides a stable physiological pH without CO_2_ supplementation, as it is buffered with non-bicarbonate ions.

Pharmaceutical inhibitors were purchased from Selleckchem: tenovin-1 (#S8000), pifithrin-□ (#S2929), Rigosertib (#S1362) and KU-55933 (#S1092).

### Viability experiments

Cells were grown in DMEM with 10% fetal bovine serum in the presence or absence of puromycin (2μg/mL). Energy depletion was achieved in cells grown in Leibovitz’s L-15 medium without glucose supplementation and hypoxia obtained by applying a 100% N2 atmosphere (ref 26700 from Air products) in an airtight chamber (ref 27310 and 27311 from Stemcell) during 24h, according to the manufacturer’s instructions. The control cells were grown in Leibovitz’s L-15 medium with 4.5 g/dL glucose supplementation under ambient atmosphere. Flow cytometry analysis for assessment of dead cells was performed with the fixable viability dye Viobility 405/452 Fixable Dye (Miltenyi Biotec^®^). Viobility dyes react with amine groups of proteins and are non-toxic. In dead cells, intracellular proteins are stained, resulting in an increase in fluorescence.

### ATP/ADP ratio measurements

ATP and ADP measurements were performed with the ApoSENSOR ADP/ATP ratio assay (Enzo Life Sciences), in accordance with the manufacturer’s instructions.

PercevalHR was used to study the ATP/ADP ratio in live individual cells in real-time experiments, spanning multiple cell cycles. Using lentiviral transformation, we generated proximal tubular cells with stable expression of PercevalHR as well as pHRed, in order to adjust the PercevalHR signal on pH variations(*12*). ATP/ADP ratio and pH were assessed by dual signal acquisition for PercevalHR (ex 436/495; em 540) and pHRed (ex 436/550; em 640), followed by ratiometric normalization using the Ratio Plus plugin in Fiji(*64*). PercevalHR (ATP/ADP ratio) was further corrected for pH by normalization on pHRed (*12*), using the Ratio Plus plugin.

### Bioinformatic analysis

#### ATP/ADP ratio

The Fiji Trackmate plugin was used to monitor single-cell ATP/ADP ratio over time, and to determine the time of cytokinesis for each cell(*65*). Briefly, every cell was assigned to a single trajectory at a specific timepoint, with a corresponding ATP/ADP value. The times of cytokinesis were noted for each cell trajectory. ATP/ADP values were represented on the y-axis as mean+/-standard error, and time was represented on the x-axis in hours from either the start of the experiment or from the time of cytokinesis. The full script is available as Supplemental data.

#### Computation of proliferation index, overall OXPHOS index, nuclear and mitochondrial OXPHOS index in bulk RNAseq data

We used a previously published and validated method to compute a proliferation index as the median value of a list of proliferation associated genes(*28*). Similarly, we computed the overall OXPHOS index as the median value of the genes listed in the human OXPHOS pathway of the (Kyoto Encyclopedia of Genes and Genomes database(*29*). We performed the same computation on the subset of mitochondria-encoded genes to obtain the mitochondrial OXPHOS index.

#### Computation of proliferation index, overall OXPHOS index, nuclear and mitochondrial OXPHOS index in single-cell RNAseq data

We used the human rejecting kidney data set (*31*) and used the Seurat package from R (*66, 67*) to check the quality of the data and to clusterize it into different cell types. Based on the top genes discriminating the different clusters (Fig. 5), we were able to identify the different intra-renal cell types. We extracted the epithelial cells and computed indices for proliferation, mitochondrial encoded OXPHOS and nuclear encoded OXPHOS as described for bulk RNA-seq in the previous paragraph. Because the single-cell depth of sequencing is much lower than for bulk RNA-seq, we used the sum of copies within the lists of genes instead of the median value.

#### Correlation analysis of proliferation with chemosensitivity

We performed a Kendall’s test to compute the correlation (correlation coefficient k and p-value) of half maximal inhibitory concentration (IC50) value with the proliferation index for every compound tested.

#### Pathway analysis of proliferation associated genes

We performed a Kendall’s test to compute the correlation (correlation coefficient k and p-value) of every gene expression value with the proliferation index. The correlations’ p-values and coefficient were then processed with the LrPath web-based software allowing a logistic regression pathway analysis to identify the gene sets enriched in genes with proliferation-correlated expression levels. Three prespecified geneset for energy-producing pathways were analyzed: “OXidative PHOSphorylation”, “Tricarboxylic Acid Cycle”, and “glycolysis”. P-values and Odds Ratios were computed as indicators of the significance and the magnitude of the enrichment of these gene sets in genes with proliferation-correlated expression values.

### Statistical analysis

Statistical analyses were conducted using JMP 11 (SAS). Log-transformation was performed to obtain normal distribution, when necessary. Proportions, 95% confidence interval and p-values were computed using the exact binomial test. Correlations were evaluated using Kendall’s test for non-parametric values. P-values were considered significant when <0.05. Comparison between groups were considered significant when the p-value was <0.05 using a Kruskall-Wallis non-parametrical test, and secondary comparisons between pairs were considered significant when the p-value was <0.05 by Wilcoxon non parametrical test. Differences in ATP/ADP changes between treatment groups over time were assessed by analysis of variance using a standard least square model.

## Supplementary Materials

**Data File S1.** Fiji Jython script for single cell ratiometric analysis

**Movie S1.** Live PercevalHR video: scratch assay.

**Movie S2.** Live PercevalHR video: effect of various cell-cycle acting molecules.

**Fig. S1:**
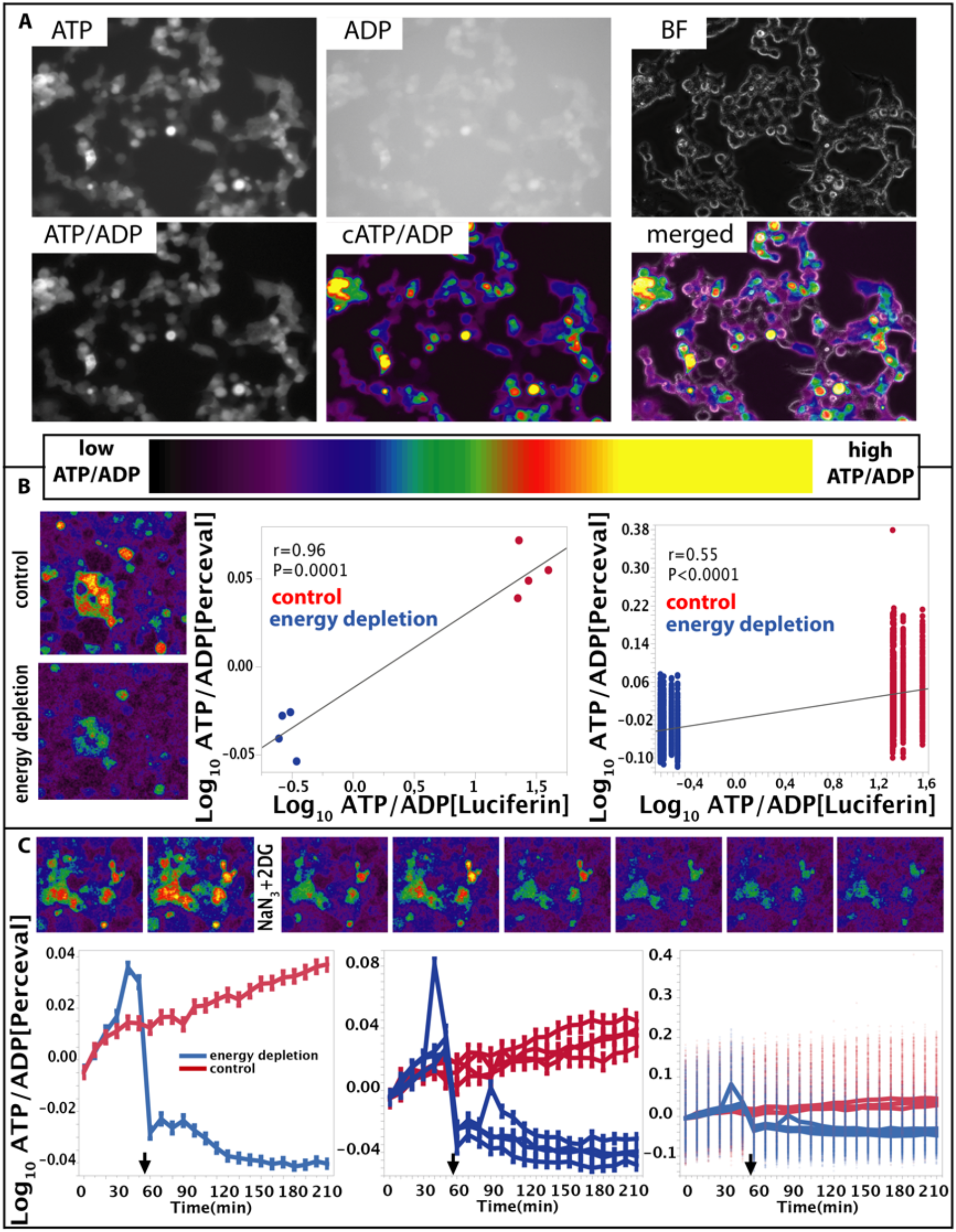
Monitoring of intracellular ATP/ADP ratio. **A.** Fluorescence imaging of ATP-bound PercevalHR (ATP), ADP-bound PercevalHR (ADP), brightfield image (BF), 2D-representation of the ATP/ADP ratio in black and white (ATP/ADP) and using artificial colors (cATP/ADP), and the cATP/ADP image merged with the brightfield image (merged). **B**. Comparison of the PercevalHR and Luciferin based assay for ATP/ADP quantification. **C**. Real time single cell ATP/ADP monitoring with PercevalHR. Left: each line represents the mean ATP/ADP value of cells from each condition (control vs energy depletion). Middle: each line represents the mean ATP/ADP value of the cells from a single well. Right: ATP/ADP values of individual cells represented by dots. The arrow indicates the time when the cells were treated with 2DG and NaN_3_ for energy depletion.

**Fig. S2.**
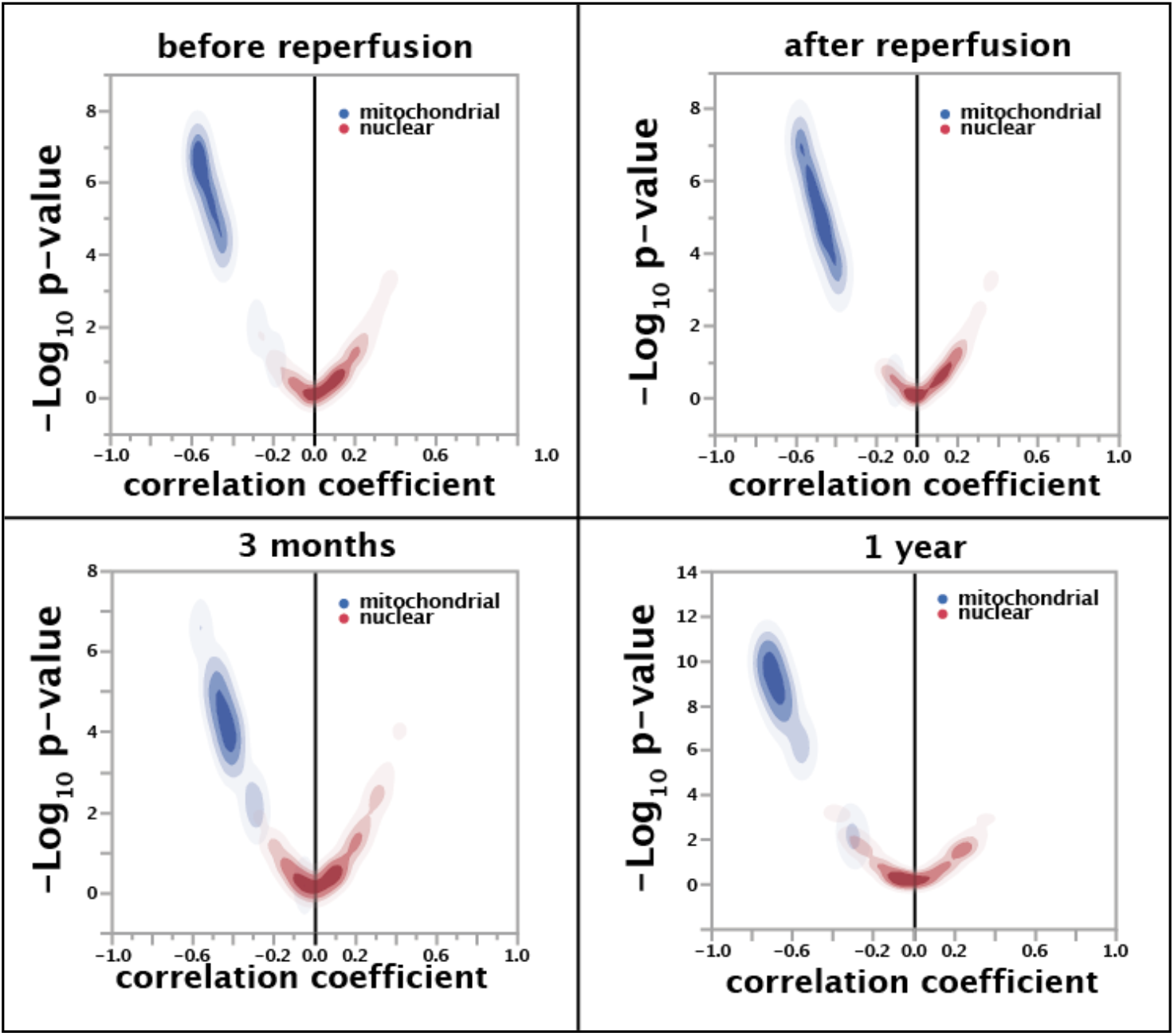
Transcriptomic signature associated with proliferation at different time points of kidney transplantation. Volcano plot showing the significance of the correlation of the proliferation index with every expression value of the OXPHOS genes at the different time points after transplantation. Nucleus-encoded genes are in red and mitochondria-encoded genes are in red.

**Fig. S3.**
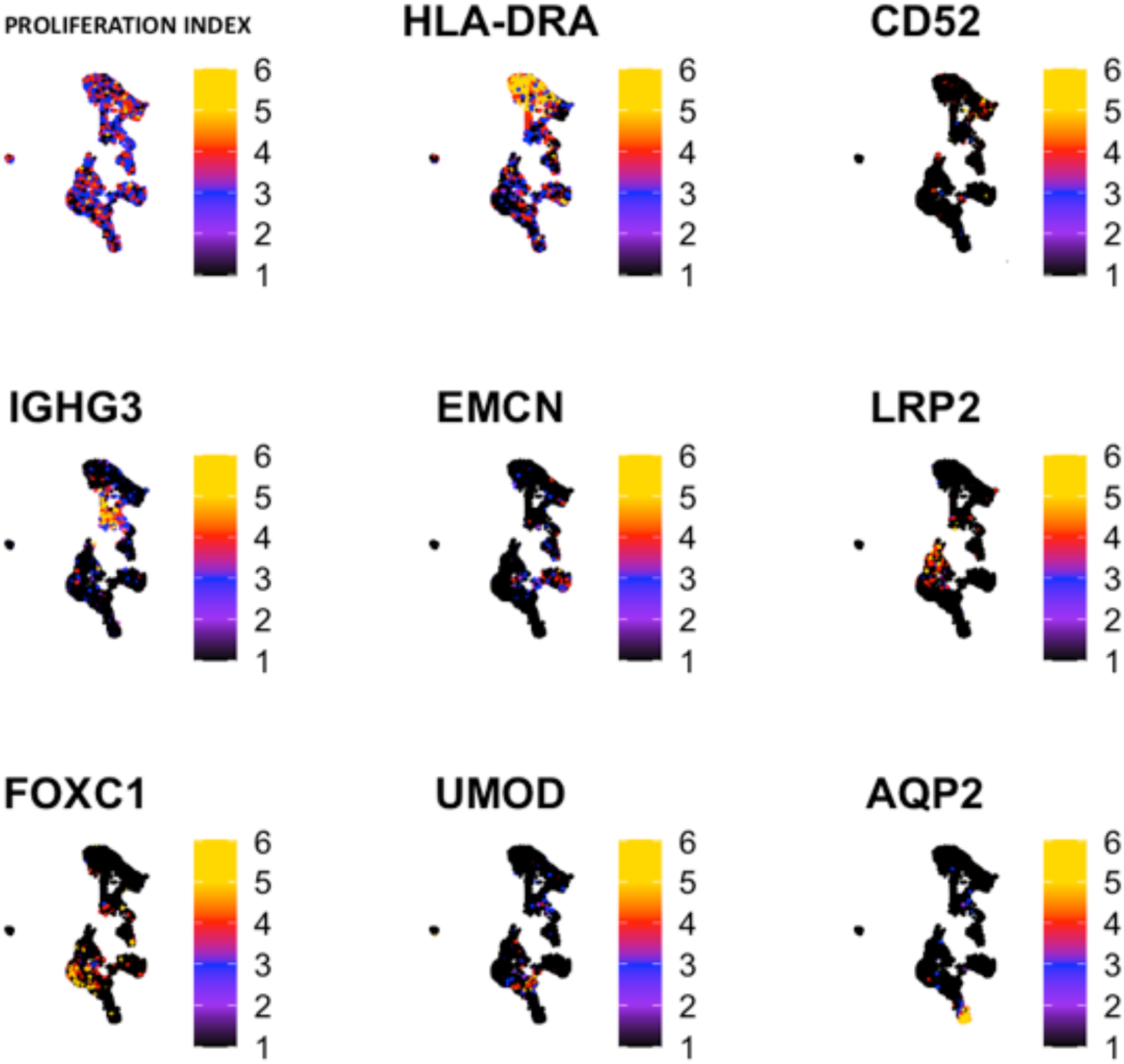
Proliferation index and renal epithelial cell differentiation markers in single cells from a rejecting human kidney. HLA-DRA: monocytes, CD52: lymphocytes, IGHG3: plasmocytes, EMCN: endothelial cells, LRP2 proximal tubule, FOXC1: descending limb of the loop of Henle, UMOD: ascending limb of the loop of Henle, AQP2: collecting duct.

**Fig. S4:**
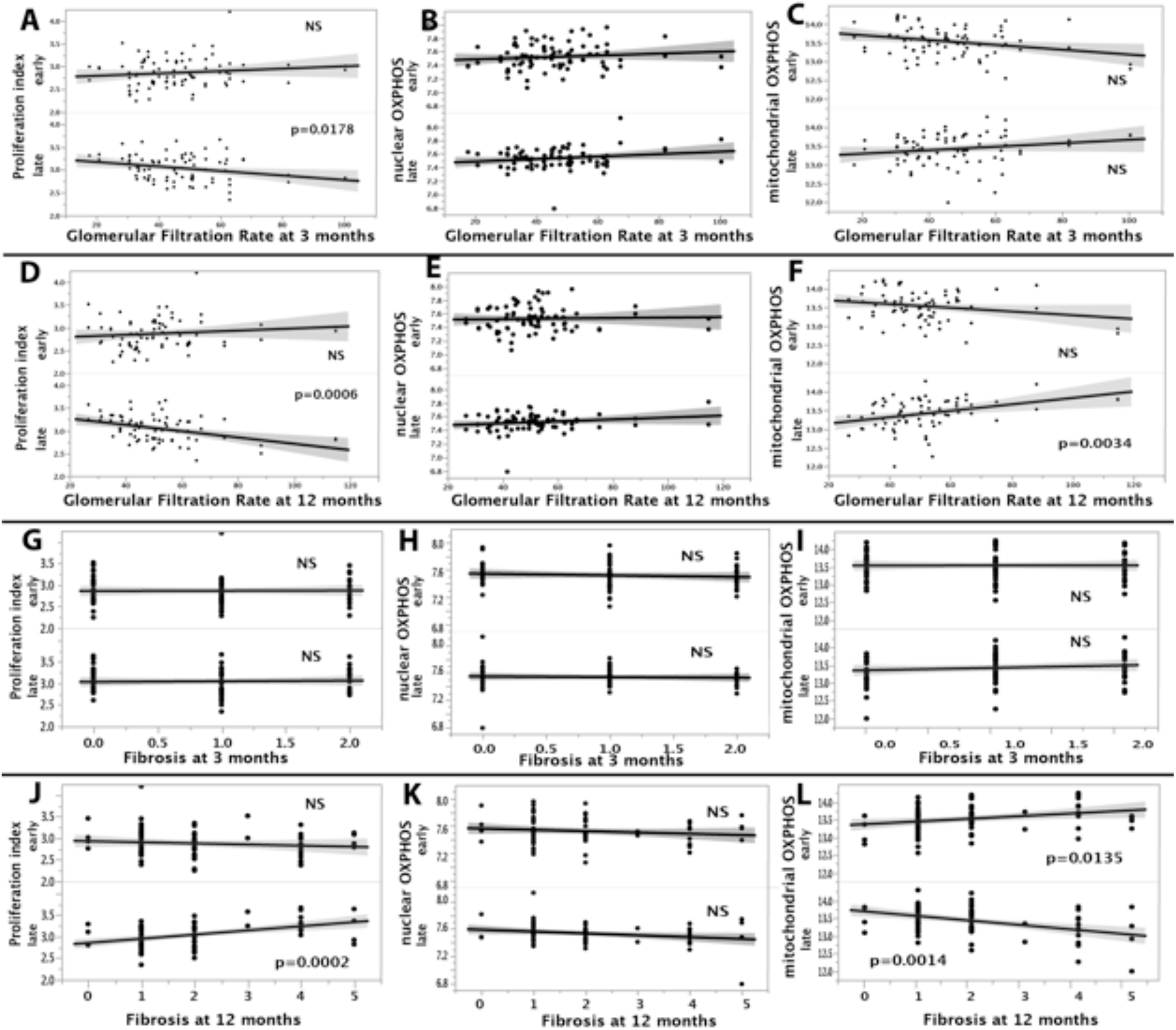
Association of the proliferation index and the OXPHOS indexes with long term outcomes. Proliferation index, overall OXPHOS index and mitochondrial OXPHOS index at acute (before transplantation and after reperfusion) and chronic time points (at 3 and 12 months after transplantation) differential correlations with 3 and 12 months glomerular filtration rate (GFR) and with 3 and 12 months kidney fibrosis, and the differential correlations of overall and mitochondrial OXPHOS with the proliferation index (p-values calculated by Kendall correlation test for non-parametric values).

**Table S1.** Top 20 Drugs with the highest correlation of chemosensitivity with proliferation, with their molecular targets and biological pathways. In blue, drugs that are more toxic in proliferating cells, and in red, drugs that are less toxic in proliferating cells. MTP: mitochondrial transition pore.

| drug (proliferation correlation<br>with chemosensitivity) | target | pathway |
| --- | --- | --- |
| NPK76-II-72-1 (+) | PLK3 | Cell cycle |
| venotoclax (+) | Bcl2 | apoptose, MTP inhibitor |
| methotrexate (+) | antimetabolite | DNA replication |
| vorinostat (+) | HDACi | histone acetylation |
| ACY-1215 (+) | HDAC6i | histone acetylation |
| navitoclax (+) | Bcl2 | apoptosis |
| daporinad (+) | NAMPT | metabolism |
| PHA-793887 (+) | CDK2/7/5 | Cell cycle |
| Ara-G (+) | antimetabolite | DNA synthesis |
| tivozanib (+) | VEGFR1/2/3 | RTK signalling |
| GSK1070916 (+) | AURKA/C | mitose |
| brivanib, BMS-540215 (+) | VEGFR/PDGFR | RTK signalling |
| kb NB 142-70 (+) | PKD | other |
| TL-2-105 (+) | CRAF inhibitor | ERK/MAPK |
| XMD13-2 (+) | RIPK1 | apoptosis |
| T0901317 (+) | LXR,FXR | other |
| alisertib (+) | AURKA | mitose |
| trametinib (-) | MEK1/2 | ERK/MAPK |
| refametinib (-) | MEK1/2 | ERK/MAPK |
| selumetinib (-) | MEK1/2 | ERK/MAPK |

**Table S2.**
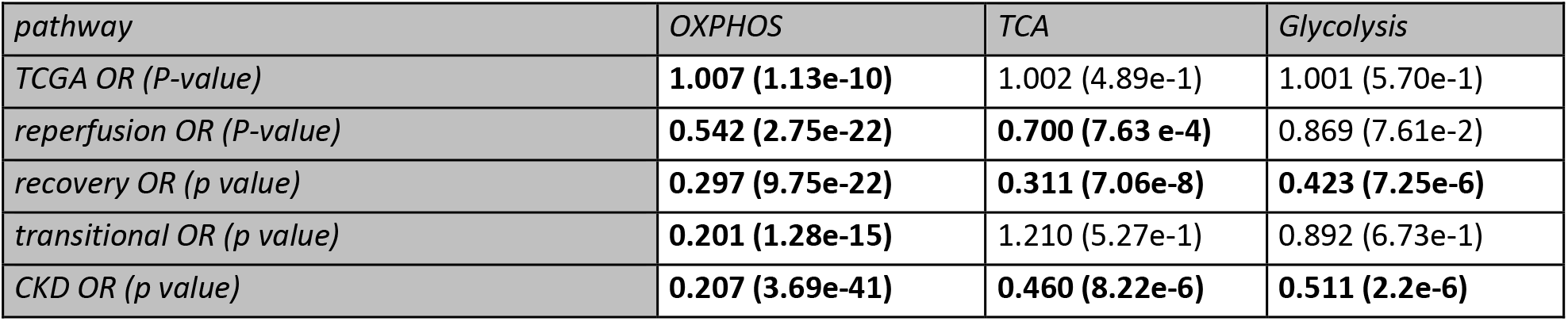
Association of proliferation-associated genes with energy metabolism pathways from the KEGG encyclopedia. The analysis was performed on cancer samples from the TCGA repository and on kidney allografts early (reperfusion) biospies or late (3 to 12 months) biopsies classified as either recovery, transitional or chronic disease. OR: Odds Ratio for genes which expression values are correlated with the proliferation index to belong to the tested pathways (versus non correlated genes).

**Table S3.**
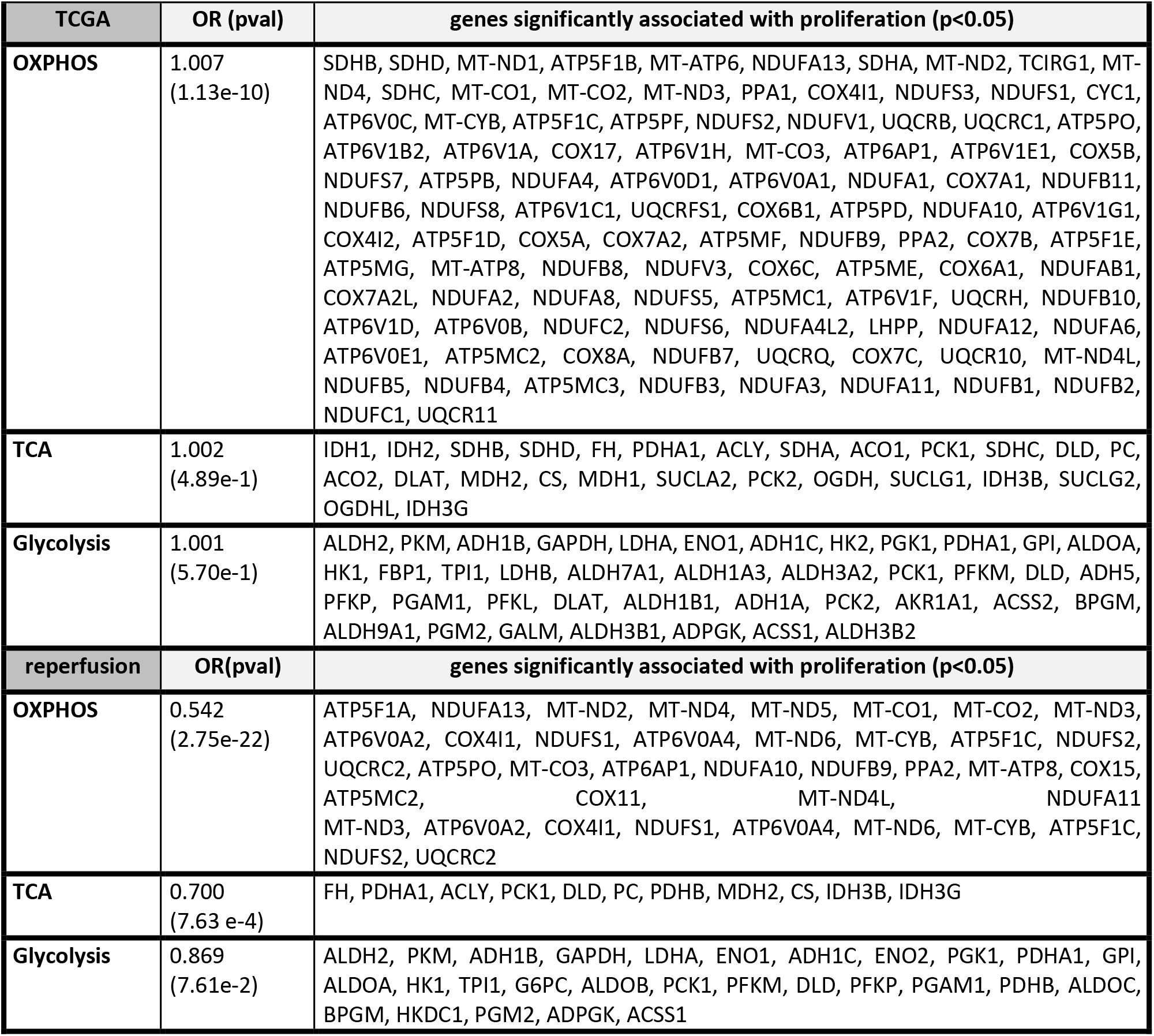
Metabolic pathways and genes associated with proliferation in tumors (TCGA) and in injured kidneys (reperfusion). OXPHOS: oxidative phosphorylation pathway; TCA: tricarboxyclic acid cycle, OR: Odds Ratio for genes which expression values are correlated with the proliferation index to belong to the tested pathways (versus non correlated genes).

**Table S4.** Key resources.

| REAGENT or RESOURCE | SOURCE | IDENTIFIER |
| --- | --- | --- |
| **Chemicals, Peptides, and Recombinant Proteins** |  |  |
| puromycin | Sigma Aldrich | P8833 |
| 4.5 g/dL glucose | Sigma Aldrich | G7021 |
| pifithrin-α | Selleckchem | S2929 |
| ku-55933 | Selleckchem | S1092 |
| rigosertib | Selleckchem | S1362 |
| tenovin-1 | Selleckchem | S8000 |
| Viability 405/452 Fixable Dye | Miltenyi Biotec® | 130-109-816 |
| **Critical Commercial Assays** |  |  |
| ApoSENSOR ADP/ATP ratio assay | Enzo | ALX-850-248-KI01 |
| **Experimental Models: Cell Lines** |  |  |
| HK-2 | ATCC | HCRL-2190 |
| **Cell culture and microscopy** |  |  |
| Leibovitz's L-15 medium with no phenol red | Fisher | 21083027 |
| airtight chamber | Stemcell | 27310 and 27311 |
| 100% N2 atmosphere | Air products | 26700 |
| **Filter sets for microscopy** |  |  |
| PercevalHR filterset | Chroma | ET436/20x, ET495/10x, T510lpxru, ET 540/40m |
| pHRed filterset | Chroma | ET550/20x, T750lpxr, ET585/20m, ET640/20m |
| **Recombinant DNA** |  |  |
| PercevalHR | Addgene | RRID:Addgene_49083 |
| pHRed | Addgene | RRID:Addgene_65742 |
| **Datasets** |  |  |
| GDSC transcriptomic and chemosensitivity data | Genomics of Drug Sensitivity in Cancer |  |
| TCGA transcriptomic data | Genomic Data Commons Data Portal |  |
| RNAseq of kidney allografts | NCBI GEO | GSE126805 |
| single cell RNAseq of rejecting allograft | NCBI GEO | GSE109564 |
| **Software and Algorithms** |  |  |
| Single cell ATP/ADP ratio script | Data File S1 | script for single cell ratiometric analysis |

## Acknowledgments

The results shown here are in part based upon data generated by the TCGA Research Network: https://www.cancer.gov/tcga.

## Funding

Institut National de la Santé et de la la Recherche Médicale ATIP Avenir program (PG)

Monahan Foundation (PG)

Fondation pour la Recherche Médicale (PG)

Groupe Pasteur Mutualité (PG)

Société Francophone de Transplantation (PG)

Société Francophone de Néphrologie Dialyse et Transplantation (PG)

Arthur Sachs fellowship (PG)

Philippe Foundation (PG)

Fulbright Scholarship (PG)

Fondation de l’Avenir (PG)

National Institute of Health/National Institute of Diabetes and Digestive and Kidney

Diseases 2R01DK072381 (JVB)

National Institute of Health/ National Institute of Diabetes and Digestive and Kidney

Diseases R37DK039773(JVB)

National Institute of Health/ National Institute of Diabetes and Digestive and Kidney

Diseases UH3 TR002155 (JVB)

## Author contributions

Conceptualization: PG

Methodology: PG, ML, LL, SV, MTV, JH, JVB

Software: PG

Investigation: PG, ML, LL, SV, DL

Resources: PG, MTV, JVB

Visualization: PG, ML, DL, JH, JVB

Funding acquisition: PG, JVB

Supervision: PG, JVB

Writing – original draft: PG, JH

Writing – review & editing: PG, ML, LL, SV, DL, MTV, JH, JVB

## Data and materials availability

Key resources are summarized in Table S4. Further information and requests for resources and reagents should be directed to and will be fulfilled by the Lead Contact, Pierre Galichon. Cell lines generated in this study are available through a Material Transfer Agreement on request. The published article includes all codes generated during this study.

